# A distinct class of eukaryotic demethylases safeguards the positional fidelity of N^6^-methyladenine (6mA)

**DOI:** 10.1101/2025.06.10.658764

**Authors:** Junhua Niu, Haoze Yu, Yongqiang Liu, Lanheng Nie, Bei Nan, Wenxin Zhang, Ni Song, Shan Gao

## Abstract

DNA N^6^-methyladenine (6mA) has recently been rediscovered as an epigenetic mark in eukaryotes. Several demethylases responsible for 6mA removal have been identified in multicellular eukaryotes. However, despite the much higher abundance of 6mA in unicellular species, no such enzymes have been reported and the biological significance of 6mA removal in these organisms remain unclear. Here, we identify and functionally characterize DMT3 as a dedicated 6mA demethylase in unicellular eukaryotes. DMT3 targets both fully and hemi-methylated ApT dinucleotides on DNA. It is predominantly localized at transcription start sites (TSS), where it antagonizes the activity of the 6mA methyltransferase AMT1. Deletion or catalytic inactivation of *DMT3* leads to genome-wide ectopic 6mA deposition specifically at TSS regions, driven by AMT1, and is accompanied by increased chromatin accessibility and elevated enrichment of transcription-associated histone marks. These epigenetic alterations disrupt gene expression, reduce osmotic stress resistance, and impair the initiation of sexual reproduction. Together, our findings uncover a critical 6mA demethylase in unicellular eukaryotes, and highlight the essential role of active 6mA removal in maintaining transcriptional and developmental integrity.

## Introduction

N^6^-methyladenine (6mA) is rediscovered as an important form of DNA methylation in eukaryotes. It has been implicated in multiple biological processes such as stress response, embryogenesis, immunity, as well as in pathological circumstances (Greer et al. 2015; Zhang et al. 2015; Wu et al. 2016; Xie et al. 2018; Kweon et al. 2019; Li et al. 2020; Zhang et al. 2020b; Sheng et al. 2021; Liu et al. 2024; Duan et al. 2025). Uncovering the enzymatic system that regulates 6mA dynamics and dictates its functions is essential for 6mA biology.

6mA in multicellular eukaryotes, especially in mammalian systems, has been intensively studied while remaining controversial. On the one hand, 6mA levels in certain systems are overestimated, possibly due to contaminations or systematic biases, despite the use of otherwise orthogonal detection methods (Douvlataniotis et al. 2020; Kong et al. 2022; Boulet et al. 2023; Feng and He 2023). There is growing consensus that 6mA in most multicellular eukaryotes may arise from RNA N^6^- methyladenosine (m6A) through the RNA salvage pathway (Bochtler and Fernandes 2020; Musheev et al. 2020). On the other hand, despite the reports of several 6mA demethylases (Greer et al. 2015; Zhou et al. 2018; Zhang et al. 2020a; Hahn et al. 2024), no 6mA methyltransferase (MTase) has been conclusively validated in multicellular eukaryotes (Greer et al. 2015; Xiao et al. 2018; Kweon et al. 2019; Woodcock et al. 2019; Luo et al. 2022; Ma et al. 2023). Moreover, 6mA sites characterized in these species rarely occur at palindromic motifs such as ApT dinucleotides (Greer et al. 2015; Zhang et al. 2015; Liang et al. 2018; Xiao et al. 2018), raising questions about their potential for stable maintenance and epigenetic inheritance.

Compared to its multicellular counterparts, 6mA in unicellular eukaryotes is introduced and maintained enzymatically. For instance, in the model ciliate *Tetrahymena thermophila*, the first eukaryote reported to have 6mA in its genomic DNA, 6mA is *de novo* established and maintained by MT-A70 family proteins, namely AMT2 with AMT5 (Cheng et al. 2025), and AMT1 with AMT6 or AMT7 (Beh et al. 2019; Wang et al. 2019; Wang et al. 2025), respectively. Notably, 6mA in unicellular eukaryotes occurs at symmetric ApT dinucleotides (Fu et al. 2015; Wang et al. 2017; Beh et al. 2019; Pan et al. 2023; Lax et al. 2024; Liu et al. 2024; Lax et al. 2025) and is transmitted semi-conservatively (Wang et al. 2019; Sheng et al. 2024), remarkably analogous to how 5mC is maintained at CpG dinucleotides (Yoder et al. 1997).

To meet the standard as a *bona fide* epigenetic mark, 6mA must be not only inheritable but also regulatable. It is therefore essential to identify and characterize enzymes that can catalyze the removal of 6mA and regulate its dynamics. Several demethylases have been identified in multicellular eukaryotes, including ALKBH1 and ALKBH4 in mammals (Kweon et al. 2019; Zhang et al. 2020a), NMAD-1 and ALKB-1 in the nematode *Caenorhabditis elegans* (Greer et al. 2015; Hahn et al. 2024), BmNMAD in silkworm (Wang et al. 2018), Homologs of ALKBH1 in plants (Zhou et al. 2018; Li et al. 2024), DMAD in the fruit fly *Drosophila melanogaster* (Zhang et al. 2015), and CcTET in the multicellular fungi *Coprinopsis cinereal* (Mu et al. 2022; Zhang et al. 2024).However, in unicellular eukaryotes where 6mA is often far more abundant, the enzymatic machinery governing its removal and the associated biological functions remain largely unexplored.

In this study, we identified and characterize DMT3, as the first 6mA demethylase in unicellular eukaryote, uncovering its essential role in modulating transcription, stress response, and sexual reproduction through site-specific 6mA removal.

## Results

### DMT3 is the 6mA demethylase in *Tetrahymena*

To determine whether DMT3 is responsible for 6mA demethylation, we generated a knockout (KO) strain deleting *DMT3* in *Tetrahymena* (Figure S1A, S1B). Mass spectrometry (MS) analysis showed that global 6mA level were elevated by about 20%-30% in Δ*DMT3* cells, significantly higher than in wild type (WT) cells (Figure 1A). This elevation was confirmed by immunofluorescence (IF) staining, showing stronger 6mA signals in Δ*DMT3* cells (Figure 1B, 1C). Single Molecule, Real-Time Circular Consensus Sequencing (SMRT CCS) results (Table S1, S2) also revealed a global 6mA upregulation across the genome, rising from 0.75% in WT to 0.91% in Δ*DMT3* cells (Figure 1D, Table S1). This increase was evident at the gene level, with higher 6mA levels observed in Δ*DMT3* cells (Figure 1E). At the level of individual 6mA sites, a substantial number of uniquely methylated sites emerged in Δ*DMT3* cells (282,562/1,017,930 =27.76% of Δ*DMT3* 6mA) (Figure 1D). Notably, 6mA penetrance (defined as the ratio between the number of 6mA sites and all adenine sites) of some originally low-methylated or unmethylated sites was dramatically increased after *DMT3* KO (Figure 1F). Moreover, depletion of *DMT3* did not increase RNA m6A levels (Figure 1G), suggesting that m6A is not a substrate of DMT3.

**Figure 1.**
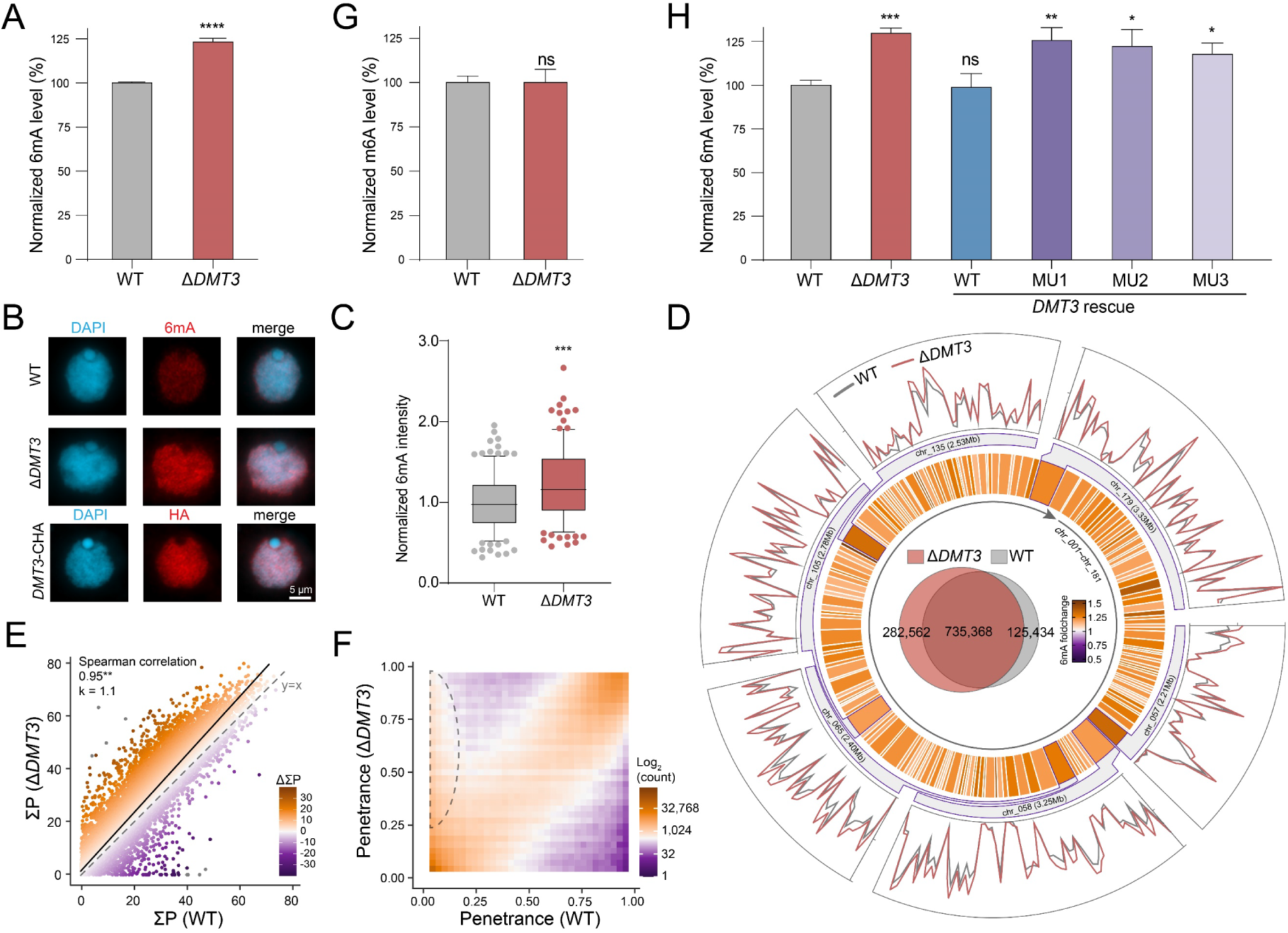
DMT3 is a potential 6mA demethylase in *Tetrahymena*. A. Mass spectrometry analysis of 6mA levels in WT and Δ*DMT3* cells. Student’s *t*-test was performed. ns *P* > 0.05, **** *P* < 0.0001. B. Immunofluorescence staining of 6mA and HA-tagged DMT3. Scale bar = 5 µm. C. Statistical analysis of 6mA signal intensity in B. Cell images were randomly selected (n = 100) and processed by ImageJ. Boxplots indicate the median, first, and third quartile. Whiskers are drawn down to the 10th percentile and up to the 90th. Points above and below whiskers are drawn individually. *** *P* < 0.001. D. Genome-wide distribution of 6mA in WT and Δ*DMT3* cells. The inner venn diagram illustrated that there are more uniquely methylated 6mA sites (6mA coverage ≥ 5×) in Δ*DMT3* cells compared to WT cells (representation factor=67.4, *P* < 0.001). The inner heatmap track showed the fold change of 6mA levels between WT and Δ*DMT3* cells. The outer line tracks showed the 6mA ratios (6mA/A) in the six longest macronuclear chromosomes of WT (blue) and Δ*DMT3* (pink) cells. E. 6mA levels of individual genes in WT and Δ*DMT3* cells. 6mA level of a specific gene was calculated as “the sum of penetrance of all 6mA sites from 100bp upstream of the TSS to 1,200bp downstream of the TES (ΣP)”. ΔΣP was calculated as ΣP_Δ_*_DMT3_* - ΣP_WT_. The Spearman’s rank correlation coefficient is significant (** *P* < 0.01). F. Density plot of 6mA penetrance (ratio between the number of 6mA and all adenines in all SMRT CCS reads at individual 6mA sites) in WT and Δ*DMT3* cells. The dashed lines illustrated some sites with extremely low penetrance in WT cells but high penetrance in Δ*DMT3* cells. G. Mass spectrometry analysis of m6A levels in WT and Δ*DMT3* during the vegetative stage. Student’s *t*-test was performed. ns *P* > 0.05. H. Mass spectrometry analysis of 6mA levels in WT, Δ*DMT3*, and rescue strains with WT or mutant copies. Student’s *t*-test was performed. ns *P* > 0.05, * *P* < 0.05, ** *P* < 0.01, *** *P* < 0.001.

To determine whether the demethylase activity of DMT3 underpins its functions, we introduced several mutants of *DMT3* back into Δ*DMT3* cells (Figure S1C, S1D). As a control, WT DMT3 (*DMT3*-WT) was reintroduced into Δ*DMT3* cells (Figure S1C, S1D). Only *DMT3*-WT was able to restore 6mA levels, whereas the catalytic mutants failed to suppress 6mA accumulation and displayed levels comparable to those in Δ*DMT3* cells (Figure 1H).

Collectively, these results support that DMT3 is the 6mA demethylase in *Tetrahymena* and its catalytic activity is required for 6mA removal.

### DMT3 can demethylate both fully and hemi-methylated 6mApT

Despite the global increase observed in Δ*DMT3* cells (Figure 1), 6mA modifications remained exclusively associated with the self-complementary ApT dinucleotide (Figure 2A, 2B). Importantly, the proportion of fully-methylated sites remained nearly unchanged (88.33% in Δ*DMT3* vs. 88.81% in WT) (Sheng et al. 2024) (Figure 2C, 2D, Table S1). This stable ratio after DMT3 KO suggests that DMT3 does not exhibit substrate preference between fully and hemi-methylated 6mApT sites. To directly test this, we overexpressed DMT3 in WT cells (Figure S1F, S1G) and detected an approximately 15% reduction in 6mA levels (Figure 2E), indicating that DMT3 can utilize fully-methylated 6mApT as substrates. Meanwhile, in Δ*AMT1* cells wherein the maintenance MTase is deleted and >95% 6mApT sites are hemi-methylated (Sheng et al. 2024), *DMT3* overexpression (Figure S1A, S1F, S1G) also led to a significant decrease in 6mA levels (Figure 2E), demonstrating that DMT3 is capable of demethylating hemi-methylated 6mApT substrates. Based on these findings, we conclude that DMT3 can act on both fully-methylated and hemi-methylated 6mApT.

**Figure 2.**
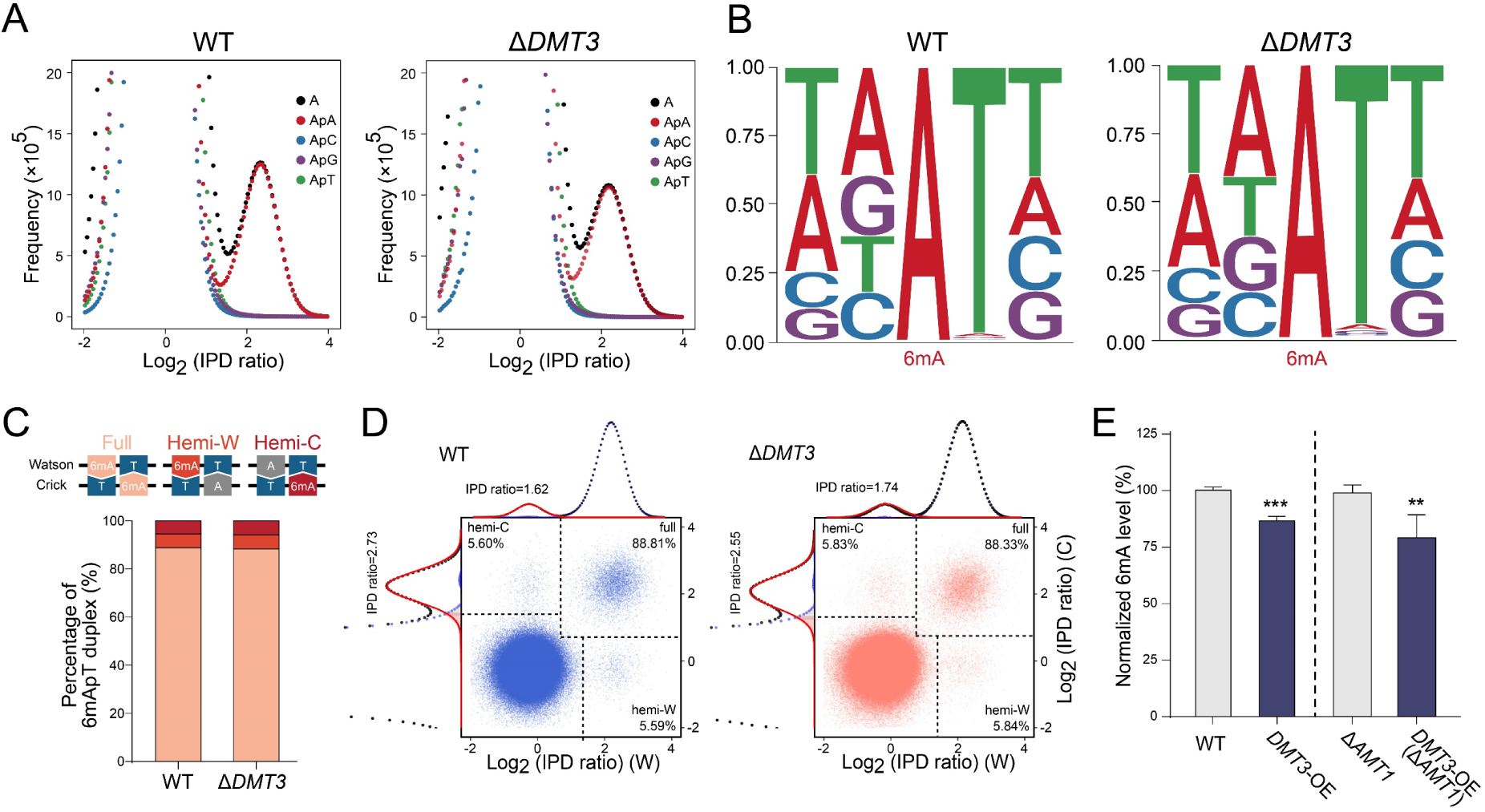
DMT3 can demethylate both fully and hemi-methylated 6mApT. A. The distribution of IPD ratio was shown for all A sites and A sites at the ApA, ApC, ApG and ApT dinucleotides, respectively. Only IPD ratios of ApT showed a bimodal distribution. B. The sequence logos for 6mA sites, revealing AT dinucleotide as the conserved motif in both WT and Δ*DMT3* cells. C. Proportion of full, hemi-W and hemi-C 6mA sites at ApT dinucleotides in WT and Δ*DMT3* cells. D. IPD ratios of each ApT dinucleotide on Watson and Crick strands in WT and Δ*DMT3* cells. Different IPD ratio thresholds were used to determine the methylation status of each ApT dinucleotide. E. Mass spectrometry analysis of 6mA levels in WT, *DMT3*-OE, Δ*AMT1*, and *DMT3*- OE (Δ*AMT1*) cells. Student’s *t*-test was performed. ** *P* < 0.01, *** *P* < 0.001.

### DMT3 ensures positional fidelity of 6mA by preventing ectopic accumulation

Genome feature annotation revealed that the increased 6mA in Δ*DMT3* cells notably occurred on intergenic regions and 5’ untranslated regions (UTR) (Figure S3A). A finer-scale analysis around gene bodies revealed that the majority of 6mA sites in Δ*DMT3* cells still accumulate in linker DNA regions between nucleosomes downstream of the TSS, similar to WT cells (Wang et al. 2017; Wang et al. 2019; Sheng et al. 2024). However, a distinct set of ectopic 6mA sites emerged at TSS in Δ*DMT3* cells, along with slight increases at pre-existing 6mA peaks (Figure 3A, 3B). These results suggest that DMT3 is preferentially affecting 6mA distributed around TSS.

**Figure 3.**
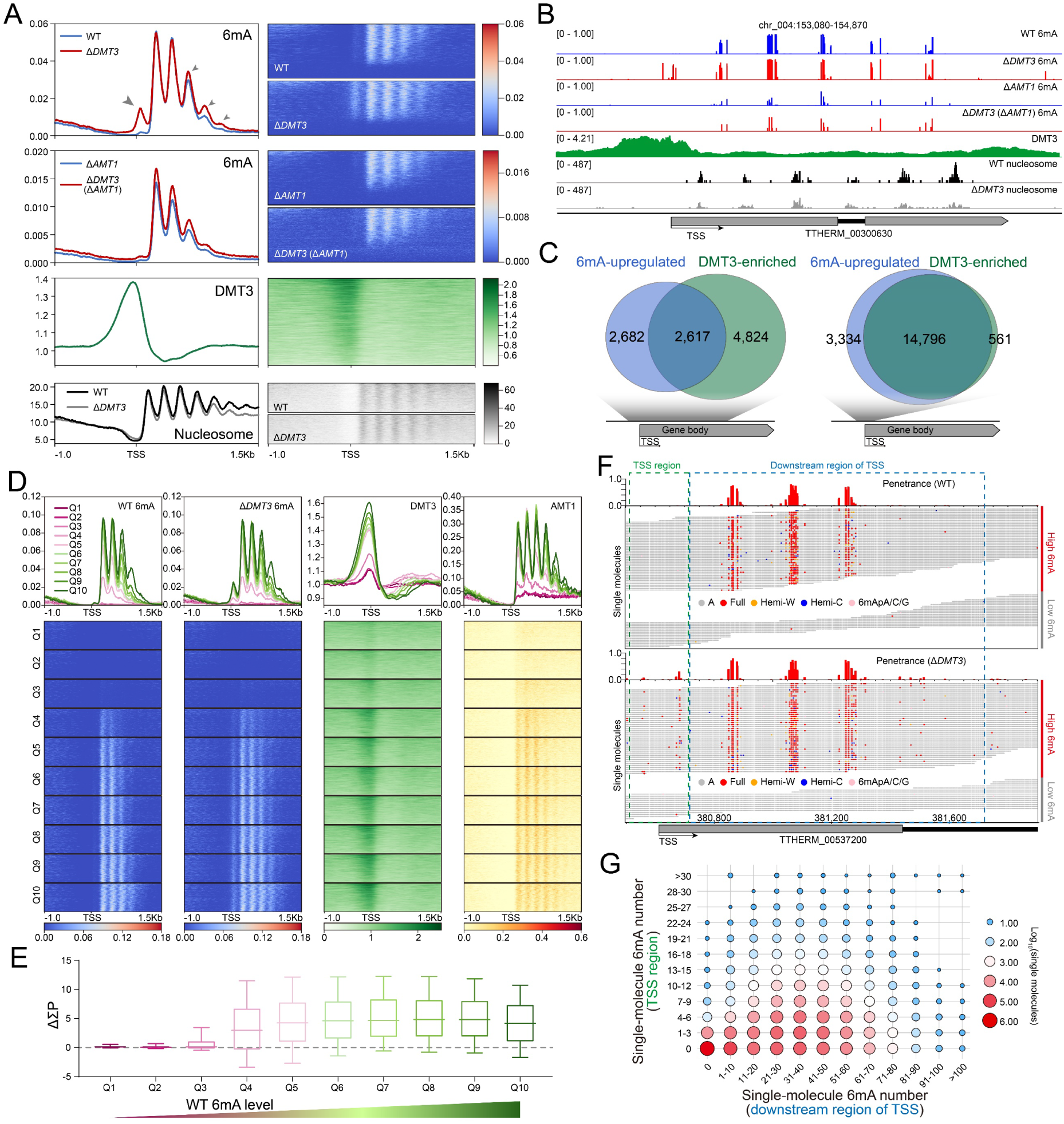
DMT3 colocalizes with the ectopic 6mA in Δ*DMT3* cells. A. Distribution profiles (left) and heatmaps (right) of 6mA in WT, Δ*DMT3*, Δ*AMT1*, and Δ*DMT3* (ΔA*MT1*) cells detected by CCS, DMT3 binding detected by ChIP-seq, and nucleosome in WT and Δ*DMT3* cells detected by MNase-seq, all centered around TSS. 6mA and nucleosome exhibited the same periodicity but in opposite phase. DMT3 was specifically enriched around the TSS. Grey arrowheads represented a newly emerged 6mA peak on TSS and elevated 6mA signal levels downstream of TSS in Δ*DMT3* cells. B. A representative genomic locus showing distribution of 6mA, DMT3, and nucleosome. C. Overlap between DMT3-enriched genes (ChIP/input ratio ≥ 1) and genes with upregulated 6mA levels (6mA foldchange ≥ 1.2) in the gene body (left, from 100 bp upstream to 1200 bp downstream of TSS, representation factor=1.5, *P* < 0.001) and around TSS (right, from 100 bp upstream to 100 bp downstream of TSS, representation factor=1.4, *P* < 0.001). Genes with extremely low 6mA levels (ΣP≤ 2.5) were excluded. D. Distribution profiles (top) and heatmaps (bottom) of 6mA, DMT3, and AMT1. Genes were ranked from low to high by their ΣP in WT cells and divided into 10 quantiles (Q1 to Q10), each containing an equal number of genes. E. 6mA level change (ΔΣP=ΣP_Δ_*_DMT3_* - ΣP_WT_) in different quantiles upon *DMT3* deletion. Gene categories were defined as in D. F. Comparison of ensemble and single-molecule level 6mA in WT and Δ*DMT3* cells, showcased by a representative genomic region. 6mA sites on each individual DNA molecules were displayed with corresponding 6mA penetrance on the top. DNA molecules were classified into high 6mA group and low 6mA group. TSS region (from 100bp upstream to 100bp downstream of TSS) was highlighted with green box. Downstream region of TSS (from 100bp downstream to 1200bp downstream of TSS) was highlighted with blue box. G. Bubble diagram showing the distribution of DNA molecules classified base on their numbers of 6mA site on TSS region and downstream region of TSS. Only DNA molecules that fully span the region from 100bp upstream to 1200bp downstream of TSS were included in the analysis

In support of its role, the endogenously tagged DMT3 was exclusively present in the macronucleus (MAC) but not in the micronucleus (MIC) (Figure 1B, S1E), consistent with the distribution of 6mA (Wang et al. 2017). More importantly, DMT3 was specifically enriched around the TSS, mirroring the position of the emerged TSS-associated 6mA peak in Δ*DMT3* cells (Figure 3A, 3B). Gene level analysis, focusing on gene bodies from 100bp upstream to 1200bp downstream of TSS, revealed a significant overlap between DMT3-enriched genes and 6mA-upregulated genes, with nearly half of 6mA-upregulated genes enriched with DMT3 (Figure 3C, left panel). When narrowing the window to 100bp upstream and 100bp downstream of the TSS, corresponding to the peak of ectopic 6mA, the overlap became even more pronounced (Figure 3C, right panel).

To further investigate the correlation between DMT3 and its targeted 6mA sites, genes were divided into ten quantiles (Q1-Q10) based on their 6mA levels in WT cells, ranging from low to high. The enrichment level of DMT3 was positively correlated with 6mA abundance in WT cells: genes with higher 6mA levels exhibited stronger DMT3 occupancy (Figure 3D). Consistently, the rise in 6mA levels in Δ*DMT3* cells was largely confined to 6mA-enriched genes, suggesting that a threshold level of 6mA is required for DMT3-mediated demethylation (Figure 3E). In support, at the single-molecule level, ectopic 6mA peaks at TSS regions in Δ*DMT3* cells occurred exclusively on DNA molecules that already carried 6mA downstream of the TSS (Figure 3F, 3G). In contrast, DNA molecules initially lacking 6mA did not acquire new 6mA marks at the TSS upon *DMT3* deletion (Figure 3F, 3G). Given that ectopic TSS-associated 6mA were mostly fully-methylated ApT (Figure 3F), these observations imply that 6mA removed by DMT3 is likely deposited by the MTase AMT1 (Wang et al. 2019). In line with this, AMT1 enrichment levels were positively correlated with 6mA abundance in WT cells (Figure 3D). More importantly, deletion of *AMT1* in Δ*DMT3* cells abolished the ectopic 6mA peak at TSS regions as expected (Figure 3A, 3B, S3C-H; Table S1), indicating that DMT3 removes 6mA deposited by AMT1 at these sites, Together, these results indicate that DMT3 does not act globally to erase 6mA, but rather functions locally to fine-tune 6mA distribution, ensuring its positional fidelity by preventing ectopic accumulation.

### DMT3-regulated 6mA modulates gene transcription and shapes the epigenetic landscape

To investigate the transcriptional consequences of *DMT3* depletion, we performed RNA-seq analysis in Δ*DMT3* cells, identifying 1,245 upregulated and 1,400 downregulated genes (Figure 4A, left). Notably, cells expressing the catalytically inactive *DMT3*-MU1 mutant exhibited transcriptional profiles highly similar to Δ*DMT3* cells (Figure 4A, right), with substantial overlap in differentially expressed genes (Figure 4B, S4A). These results suggest that the transcriptional effect of DMT3 is primarily mediated by its 6mA demethylase activity, rather than by a direct role as a transcription factor.

**Figure 4.**
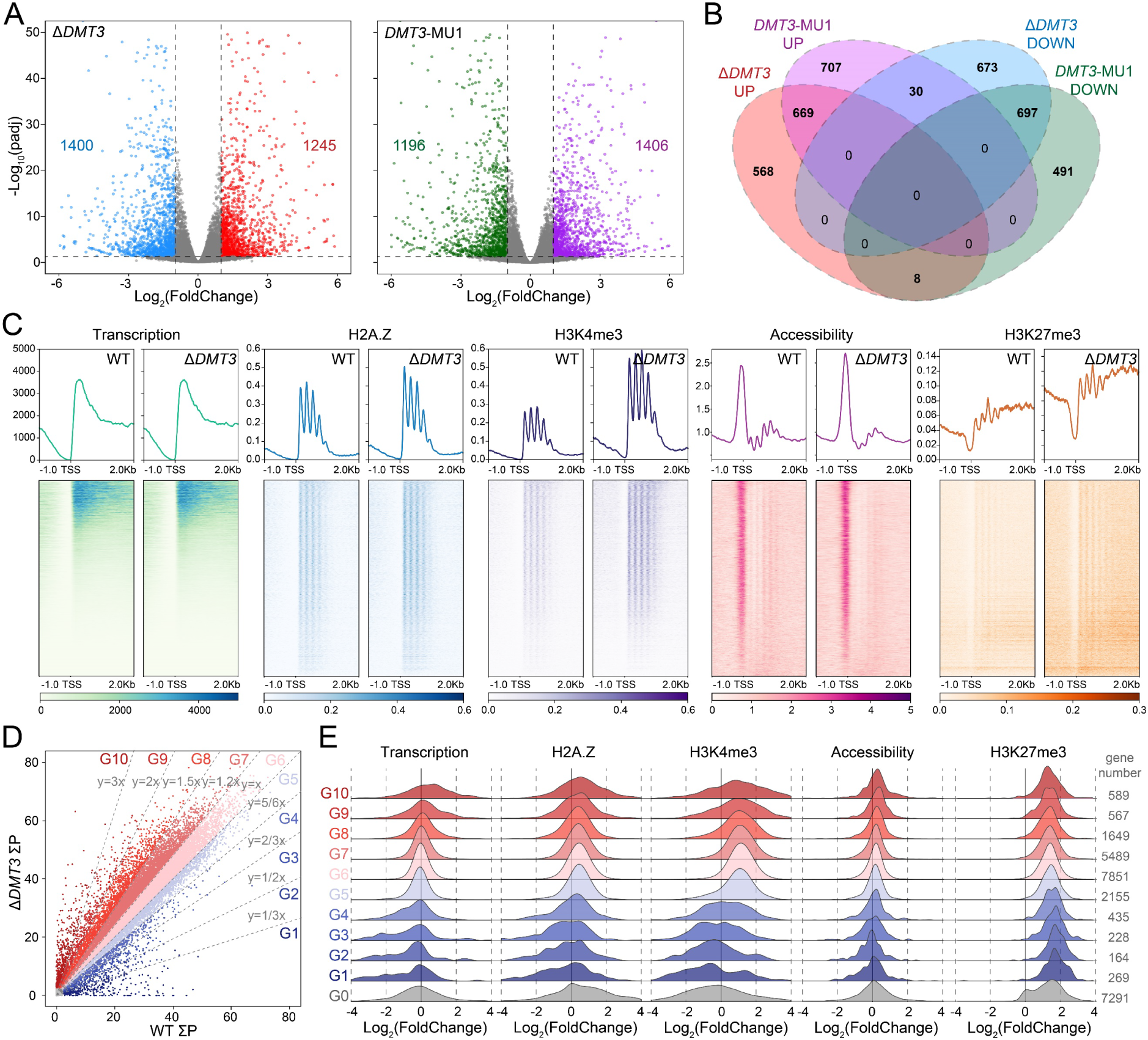
6mA is associated with transcription, H2A.Z, H3K4me3, H3K27me3, and chromatin accessibility. A. Global transcriptome analysis in Δ*DMT3* and *DMT3*-MU1 cells. Differentially expressed genes (DEGs) were highlighted. Down-regulated or up-regulated genes were defined as Log_2_(FoldChange) ≥ 1 or ≤ −1, respectively, with *P* ≤ 0.05. B. Overlap of DEGs between Δ*DMT3* and *DMT3*-MU1 cells. C. Distribution profiles (top) and heatmaps (bottom) of transcripts, H2A.Z, H3K4me3, H3K27me3, and chromatin accessibility in WT and Δ*DMT3* cells. D. 6mA levels of individual genes in WT and Δ*DMT3* cells. Genes were categorized into 10 groups (G1-G10, from low to high) based on the fold change of their ΣP (ΣPΔ*DMT3*/ΣPWT): ≤ 1/3, 1/3 - 1/2, 1/2 - 2/3, 2/3 - 5/6, 5/6 - 1, 1 - 1.2, 1.2 - 1.5, 1.5 - 2, 2 - 3, and >3. Genes with extremely low ΣP (≤ 2.5) were assigned to G0 (grey box, left bottom corner). E. Log2(FoldChange) distribution of transcription, H2A.Z, H3K4me3, H3K27me3, chromatin accessibility for each group of genes in D.

To explore the effect of 6mA perturbation on the chromatin landscape, we performed native ChIP-seq for two active transcription-associated marks, the histone modification H3K4me3 and the histone variant H2A.Z (Wang et al. 2017; Duan et al. 2025; Wang et al. 2025). We also examined chromatin accessibility by ATAC-seq. Pairwise comparison demonstrated that 6mA, transcription, H2A.Z, H3K4me3, and chromatin accessibility were strongly correlated with each other, in both WT and Δ*DMT3* cells (Figure S4C). Compared to WT cells, both H3K4me3 and H2A.Z displayed increased enrichment on nucleosome arrays downstream of the TSS in Δ*DMT3* cells, with a moderate increase for H2A.Z and a significant increase for H3K4me3 (Figure 4C). This enrichment was not due to global increases in protein levels, as Western blot analysis confirmed that their overall abundance remained unchanged (Figure S4B). Meanwhile, accessibility profiles in Δ*DMT3* cells were largely similar to those in WT cells, with a moderate increase at TSS regions (Figure 4C).

We next divided genes into 11 groups (G0-G10, from low to high) according to their fold changes in 6mA levels in Δ*DMT3* cells (Figure 4D). Groups G1 to G5 included genes with reduced 6mA, while G6 to G10 included genes with increased 6mA. Quantitative analysis revealed coordinated changes in 6mA, transcription, activating histone marks (H2A.Z and H3K4me3), and chromatin accessibility in G1-G10. These features tended to increase in groups displaying 6mA upregulation (G6-G10) and decrease in groups with 6mA downregulation (G1-G5). Among them, changes in H2A.Z and H3K4me3 aligned more closely with 6mA dynamics than did chromatin accessibility (Figure 4E).

Group G0 consisted of genes with negligible 6mA in both WT and Δ*DMT3* cells (Figure S4D), consistent with our finding that *DMT3* loss only affected 6mA-bearing genes (Figure 4D, 4E). They showed markedly higher H3K27me3 level and minimal levels of transcriptional activity, activating marks, and chromatin accessibility (Figure S4D), suggesting their expression is predominantly controlled by H3K27me3- mediated repression (Liu et al. 2007). Being already transcriptionally silent, they were largely unresponsive to further H3K27me3 accumulation in Δ*DMT3* cells (Figure 4E).

Taken together, *DMT3* knockout-induced 6mA upregulation promotes transcription, potentially through increased levels of transcription-associated epigenetic marks and enhanced chromatin accessibility.

### Defects in stress response and sexual reproduction initiation in Δ*DMT3* cells

To investigate the physiological relevance of DMT3, we first evaluated the impact of its loss on asexual growth. GO enrichment analysis of upregulated genes in Δ*DMT3* cells revealed significant enrichment in pathways associated with transmembrane transportation, likely reflecting altered osmotic regulation (Figure 5A). To assess the osmotic stress response, we compared the growth of WT and Δ*DMT3* cells. Both WT and Δ*DMT3* cells showed comparable growth in regular SPP medium (Figure 5B). However, upon NaCl supplementation, Δ*DMT3* cells showed a marked growth defect, whereas WT cells tolerated elevated salt concentration up to 180mM without significant impairment (Figure 5B). This defect in osmotic pressure response could be rescued by reintroducing WT DMT3, but not by catalytic mutants (Figure 5B), indicating that the demethylase activity of DMT3 is required for this function. For functional validation, we selected four genes associated with ion transport across the membrane from the enriched gene set with co-upregulated levels of transcription and H3K4me3 in Δ*DMT3* cells (Figure 5C). Notably, overexpression of each individual gene led to growth defect under high osmotic pressure, although to a lesser extent than in Δ*DMT3* cells (Figure 5D, S1H, S1I), suggesting that misregulation of these genes contributes to the osmotic sensitivity observed in Δ*DMT3* cells. Together, these findings support a direct involvement of DMT3-mediated 6mA demethylation in osmotic stress adaptation.

**Figure 5.**
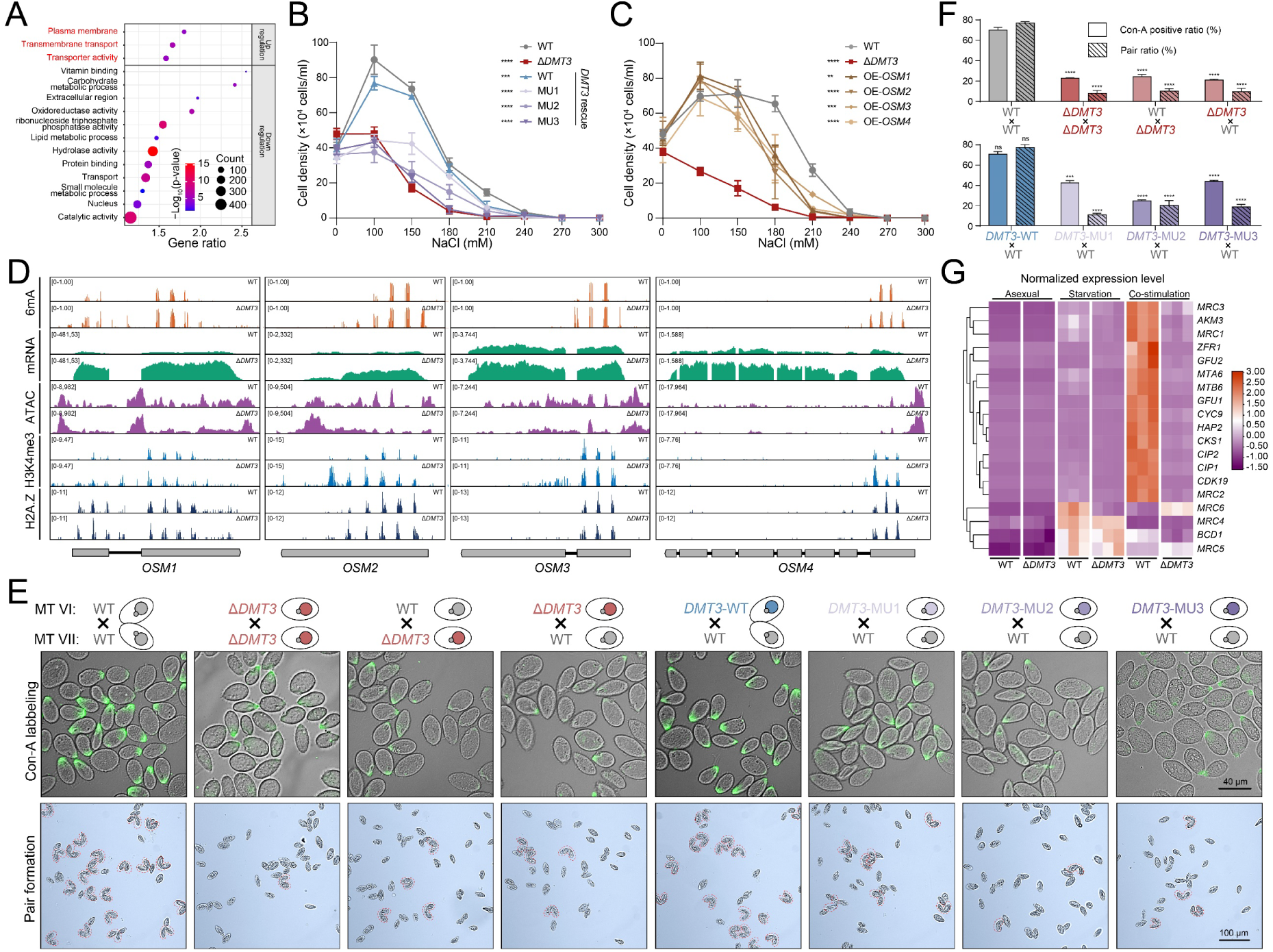
Defects of Δ*DMT3* cells in stress response and conjugation initiation. A. Gene ontology enrichment analysis of DEGs in Δ*DMT3* cells. B. Cell density curves of WT, Δ*DMT3*, and rescue cells. Cells were counted 48 hours after inoculation in SPP medium with three replicates. Two-way ANOVA: *** *P* < 0.001 and **** *P* < 0.0001. C. Cell density curves of WT, Δ*DMT3* and cells overexpressing genes related to osmotic pressure regulation, under different concentrations of NaCl in SPP medium supplemented with 0.2 μg/mL Cd^2+^ to induce gene overexpression. The overexpressed genes were clustered in G9, G8, G8, and G7, respectively, in Figure 4E. Cells were counted 48 hours after inoculation in SPP medium with three replicates. Two-way ANOVA: ** *P* < 0.01, *** *P* < 0.001, and **** *P* < 0.0001. D. Gbrowser view of representative genes related to osmotic pressure regulation in Figure 5C, showing profiles of 6mA, transcription, H2A.Z, and H3K4me3. E. Microscopy observation of Con-A labeling and pair formation in different crosses. Scale bar = 100 µm. MT: mating type. F. Statistical analysis of Con-A labeling and pair formation in Figure 5E. Experiments were repeated three times, with > 60 cells counted for each pairing. Two-way ANOVA: ns *P* > 0.5, *** *P* < 0.001, and **** *P* < 0.0001. G. Expression profiles of genes essential for pair formation in WT and Δ*DMT3* cells at asexual growing, starvation, and co-stimulation stages.

We next investigated the influence of *DMT3* loss on sexual reproduction (referred to as conjugation). After nutrient depletion, *Tetrahymena* cells of two different mating types undergoes co-stimulation, a prerequisite step for pair formation and conjugation initiation (Finley and Bruns 1980). However, the mating efficiency of Δ*DMT3* cells (Δ*DMT3* × Δ*DMT3*) was severely impaired (Figure 5E, 5F); very few cells could undergo pairing when most WT cells have already paired, even after an extended period of mixing (Figure S5A). This failure of co-stimulation was further confirmed by the absence of concanavalin A (Con-A) receptor staining at the anterior tip of cells, a typical marker of successful co-stimulation (Figure 5E, 5F) (Wolfe et al. 1986; Pagliaro and Wolfe 1987; Wolfe and Feng 1988). Notably, the mating failure was also observed in DMT3 mutants and could not be rescued by mating with WT cells (Δ*DMT3* × WT or *DMT3*-MU1/MU2/MU3 × WT) (Figure 5E, 5F), suggesting that efficient 6mA demethylation by DMT3 is intrinsically required for conjugation initiation. Consistent with the phenotypic defects, genes involved in co-stimulation (Ma et al. 2020; Ma et al. 2024; Pinello et al. 2024; Yan et al. 2024) were almost all highly expressed in WT cells but remained silenced in Δ*DMT3* cells (Figure 5G). Intriguingly, the alteration of 6mA (increased) was inversely associated with changes of gene expression (downregulated) for most co-stimulation genes (Figure S5B), suggesting the presence of upstream regulator(s) whose DMT3/6mA-dependent misregulation may underlie the broader transcriptional silence of the co-stimulation program.

## Discussion

6mA is abundant and widely distributed in unicellular eukaryotes (Fu et al. 2015; Wang et al. 2017; Beh et al. 2019; Pan et al. 2023), yet no 6mA demethylase has been identified in these organisms to date. In this study, we identify DMT3 as the first 6mA demethylase in unicellular eukaryotes. Deletion or catalytic inactivation of DMT3 leads to a global increase in 6mA levels, while its overexpression results in a marked reduction. DMT3 exhibits demethylation activity toward both fully and hemi-methylated 6mApT sites. It is enriched at TSS regions of 6mA-marked genes, where it specifically removes 6mA deposited by the MTase AMT1. Loss of DMT3 enzymatic activity alters gene transcription and reshapes global epigenetic landscape, affecting key biological processes such as osmotic response and sexual reproduction initiation. Collectively, our findings establish DMT3 as a *bona fide* 6mA demethylase in unicellular eukaryotes and underscore the functional significance of active 6mA removal in transcriptional regulation and cellular fitness (Figure 6).

**Figure 6.**
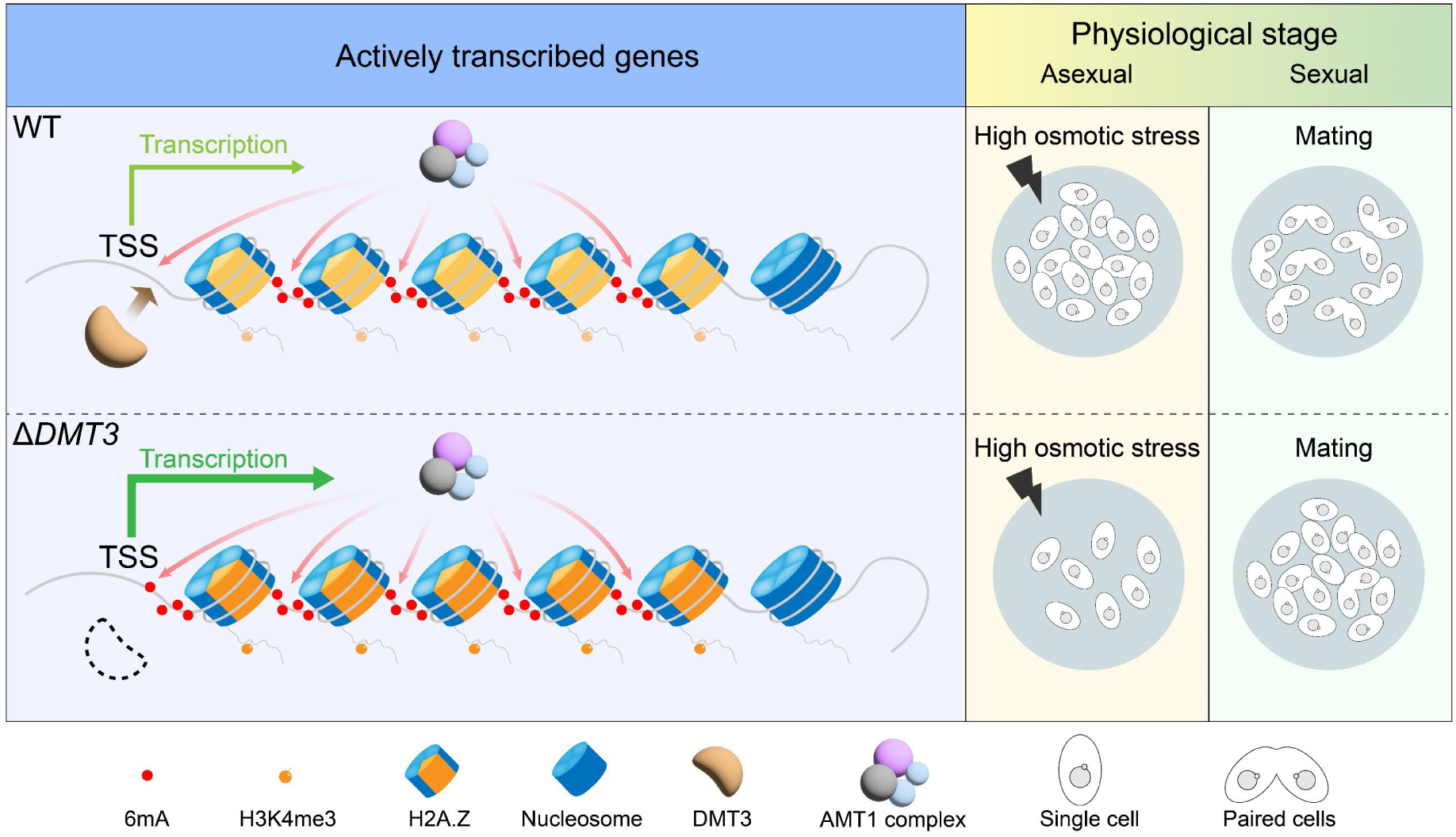
A diagram summarizing the epigenetic regulatory functions of DMT3. The tug-of-war between AMT1-mediated 6mA methylation and DMT3-mediated demethylation at transcription start sites (TSSs). In wild-type cells, a dynamic balance between methylation and demethylation is maintained during each cell cycle to keep TSSs unmethylated. Loss of DMT3 disrupts this equilibrium, leading to ectopic 6mA accumulation at TSSs and altered transcriptional and physiological responses.

Previous studies have shown that the AMT1 complex is recruited by di- nucleosomes enriched with H2A.Z and H3K4me3 and that loss of either mark on either side diminishes 6mA deposition (Wang et al. 2025). It however remains unclear why the nucleosome-free regions (NFRs) upstream of the +1 nucleosome stay unmethylated (Beh et al. 2019; Wang et al. 2019; Wang et al. 2025), despite notable enrichment of AMT1 complex components at the +1 nucleosome (Wang et al. 2025). In this study, we demonstrate that the unmethylated status of TSS was achieved by DMT3-mediated 6mA removal that counteracts AMT1 activity. Both gene-level and single-molecule analyses reveal that ectopic 6mA accumulation in *DMT3*-deficient cells preferentially occurs on DNA molecules already enriched with AMT1-deposited 6mA, suggesting a “methylation-demethylation tug-of-war” at regulatory regions. Given that AMT1 functions as the maintenance methyltransferase during each cell cycle to restore 6mA patterns by semi-conservative transmission (Sheng et al. 2024), our findings suggest that DMT3 must likewise act during each cycle to antagonize AMT1 activity specifically at TSS regions. Such dynamic, cell cycle-coupled interplay between AMT1-mediated deposition and DMT3-mediated removal constitutes a regulatory mechanism that maintains the positional specificity of 6mA and thereby preserve proper transcriptional regulation (Figure 6). This mechanism may also operate in other unicellular eukaryotes, such as the green alga *Chlamydomonas reinhardtii*, where 6mA exhibits a bimodal distribution flanking the TSS and the TSS itself remains unmethylated (Fu et al. 2015).

In multicellular eukaryotes, reported 6mA demethylases often prefer non-canonical DNA structures, such as bulged or bubbled DNA (Li et al. 2020; Zhang et al. 2020a). In contrast, our systematic knockout and rescue experiments demonstrated that DMT3 can remove both fully and hemi-methylated 6mA, supporting its activity on double- stranded (ds) DNA substrates. Structural modeling further supported this, suggesting that DMT3 engaged dsDNA by flipping the methylated base into its catalytic pocket, a mechanism reminiscent of 5mC demethylation by TET family proteins (Hu et al. 2013; Hu et al. 2015) (Figure S8). This substrate specificity aligns with the genomic distribution of 6mA in unicellular eukaryotes. Unlike in mammals, where 6mA is frequently associated with single-stranded DNA (ssDNA) regions (Li et al. 2020; Tian et al. 2020), 6mA in *Tetrahymena* and other unicellular species is predominantly enriched within gene bodies that largely adopt a double-stranded conformation with minimal presence of ssDNA features.

Compared to 5mC demethylation in higher eukaryotes, DMT3-regulated 6mA demethylation exhibits several fundamental distinctions. First, DMT3 acts broadly across the majority of 6mA-marked genes during the asexual growth, a stage analogous to somatic cell division. This widespread demethylation pattern, likely reflecting the pervasive distribution of 6mA in the *Tetrahymena* genome where over 90% of genes carry this modification (Wang et al. 2019; Sheng et al. 2024), contrasts sharply with the locus-specific nature of 5mC demethylation in somatic cells (Wang et al. 2015; Yu et al. 2015; Onodera et al. 2021; Wang et al. 2021). Such an indispensable demethylation mechanism is distinct from 5mC demethylation, which is typically triggered in response to external or internal cues (Wang et al. 2015; Yu et al. 2015; Onodera et al. 2021; Wang et al. 2021). This strategy of broad demethylase targeting and TSS protection may be conserved in certain unicellular organisms, such as green algae and ciliates, where a substantial proportion of genes exhibit high levels of 6mA and their 6mA often accumulates near TSS (Fu et al. 2015; Romero Charria et al. 2024).

Interestingly, global 6mA demethylation is not required for ciliates during gametogenesis and zygotic nucleus formation. This is because both the gametic and the zygotic nuclei originate from the parental micronucleus (MIC), which is inherently devoid of 6mA (Harrison and Karrer 1985; Karrer 2012; Zhao et al. 2021; Tian et al. 2022; Wei et al. 2022). This developmental strategy, facilitated by the ciliate-specific binucleated architecture, differs fundamentally from organisms where a single nucleus carries inherited DNA methylation that must be reset during sexual reproduction (Gu et al. 2011; Xu et al. 2011; Lee et al. 2014; Gkountela et al. 2015). However, the absence of a global 6mA erasure phase does not imply that DMT3 is dispensable during sexual development. Functional studies have shown that loss of DMT3 demethylase activity impairs conjugation competency, and transcriptomic data reveal a pronounced upregulation of *DMT3* following zygotic nucleus formation (Figure S7C). Future studies should aim to dissect the mechanisms by which DMT3-mediated 6mA demethylation contributes to nuclear differentiation in binucleated ciliates, and explore how it functions in other unicellular eukaryotes with divergent life cycles.

## Materials and Methods

### Cell culture

*Tetrahymena thermophila* wild-type strains, SB210 and CU428, were obtained from the *Tetrahymena* Stock Center (http://tetrahymena.vet.cornell.edu). Knockout, overexpression, rescue, mutation, and tagging strains used in this study were listed in Table S3. Cells were grown in SPP medium (1% protease peptone, 0.2% dextrose anhydrous, 0.1% yeast extract, 0.003% ethylenediaminetetraacetic acid (EDTA) ferric sodium salt) at 30°C.The knockout, rescue, tagging and overexpression strains were generated as described previously (Gao et al. 2013; Wang et al. 2017).

For starvation, cells at ∼2 × 10^5^ cells/mL were maintained in 10 mM Tris-HCl (pH 7.4) for at least 16 h. For conjugation, starved cells of two different mating types were mixed in equal proportions, Pair ratio were calculated and photos were taken at 6 h after mixing. For co-stimulation experiments, starved cells were mixed for only 1 hour.

For osmotic pressure test, cells with a starting concentration of ∼1 × 10^4^ cells/mL Were cultured in SPP medium containing different NaCl concentrations (0, 100, 150, 180, 210, 240, 270, 300 mM) for 48h before counting.

### Immunofluorescence staining

Cells were fixed with 2% paraformaldehyde (Sigma, P6148), as described previously (Gao et al. 2013; Wang et al. 2017). The primary antibodies were ɑ-6mA (Synaptic Systems, 202003, 1:2000, rabbit) and ɑ-HA (Cell Signaling, C29F4, 1:200, rabbit). The secondary antibodies were Goat anti-Rabbit IgG (H+L), Alexa Fluor 555 (Invitrogen, A21428, 1:4000) and goat anti-mouse IgG (H+L), Alexa Fluor 555 (Invitrogen, A28180, 1:4000). Digital images were collected using a Zeiss Axio Imager Z2 microscope with an Axiocam 506 mono camera.

### Con-A labeling

Con-A labeling was performed as previously reported (Ma et al. 2020; Yan et al. 2024). Cells were first fixed with 4% paraformaldehyde (Sigma, P6148) and washed with 0.1 M PB buffer for three times. Fixed cells were then stained with fluorescein-labeled Con-A (Vector Laboratories, Burlingame, CA, USA) at 25 μg/mL for 5 min washed with 0.1 M PB buffer for three times. Digital images were collected using a Zeiss Axio Imager Z2 microscope with an Axiocam 506 mono camera.

### Mass spectrometry analysis

Genomic DNA was treated following established protocols (Wang et al. 2019). 500 ng genomic DNA of *Tetrahymena thermophila* was digested into mononucleotides. Samples were analyzed by ultra-high-performance liquid chromatography-tandem mass spectrometry (UHPLC-QQQ-MS/MS) on an Acquity BEH C18 column (75 mm × 2.1 mm, 1.7 μm, Waters, MA, USA), using a Xevo TQ-XS triple quadrupole mass spectrometer (Waters, Milford, MA, USA) or a Hypersil GOLD column (100 x 2.1 mm, 1.9 µm, Thermo scientific) using a TSQ Quantiva mass spectrometer (Thermo scientific). The mass spectrometer was set to multiple reaction monitoring (MRM) in positive electrospray mode. For DNA 6mA and A, the selective MRM transitions were detected under m/z 266/150 and m/z 252/136, respectively. For RNA m6A and A, the selective MRM transitions were detected under m/z 282/150 and m/z 268/136, respectively. The ratio of DNA 6mA/A and RNA m6A/A was quantified by the calibration curves of nucleoside standards running at the same time.

### SMRT sequencing and data analysis

Genomic DNA was extracted using QIAGEN Genomic-tips (10223). The quality and concentration of DNA were assessed by agarose gel electrophoresis and the Qubit®3.0 Fluorometer (Thermo Fisher Scientific). SMRT sequencing data were analyzed following previously described procedures (Sheng et al. 2024; Li et al. 2025). Low-confidence single molecules (passes < 30×, standard deviation of IPD ratio in both Watson and Crick strands ≥ 0.35) were removed. After aligning CCS reads to the MAC genome (Ye et al. 2025), reads with mapped coverage less than 98% and mapped identity less than 95% were excluded.

6mA level of individual A site (penetrance) was calculated as the ratio between the number of 6mA sites and all adenine sites in all SMRT CCS reads. 6mA level of a specific gene (sigma P, ΣP) was calculated as the sum of penetrance of all 6mA sites.

### RNA sequencing and data analysis

Total RNA from three replicates of each strain was extracted using FastPure Cell/Tissue Total RNA Isolation Kit V2 (Vazyme, RC112). RNA samples were sequenced by the Illumina HiSeq platform, with 150bp pair-end sequencing (PE150). After trimming sequencing adapters and filtering out low quality reads using TrimGalore (length > 36bp and quality > 20) (Bolger et al. 2014), the numbers of reads mapped to the genome were determined using the HISAT2 software (Anders et al. 2014). Reads with multiple mapping in the SAM file were deleted. Samtools (Li et al. 2009) was used to sort and remove potential PCR duplicates. FeatureCounts (Liao et al. 2014) was implemented for counting reads to genomic features with the assembled transcripts as reference. Differentially expressed genes (DEGs) were also identified by DESeq2 (Love et al. 2014) (Log2(FoldChange) ≥ 1 or ≤ −1, P < 0.05). Gene Ontology (GO) enrichment analysis was performed using TBtools (Chen et al. 2020).

### Native ChIP-seq and data analysis

Native chromatin immunoprecipitation (ChIP) followed established protocols (Chen et al. 2016; Duan et al. 2021; Wang et al. 2025). Approximately 5 × 10^7^ purified MACs were digested by Micrococcal Nuclease (MNase, NEB) (200 U/mL, 25°C, 15min) to generate mononucleosomes. After extraction in T150 (150 mM NaCl, 0.1% Triton X- 100, 1 × Proteinase inhibitor (Roche Applied Science, 11873580001)) buffer, the solubilized chromatin was immunoprecipitated with anti-HA magnetic beads (Thermo Scientific, 88836) or protein A magnetic beads (Thermo Scientific, 10002D) preincubated with ɑ-H3K4me3 antibody (abcam, ab8580, 1:500, rabbit) at 4°C overnight. The HA-tagged protein was eluted with the HA peptide (Sigma, E6779), and the H3K4me3 enriched chromatin was eluted with SDS elution buffer (50 mM NaCl, 50 mM Tris-HCl pH 7.5, 5 mM EDTA, 1% SDS). DNA was collected by phenol-chloroform extraction for sequencing.

ChIP samples were sequenced by Illumina NovaSeq X Plus PE150. The low quality paired-end reads of raw data were filtered and sequencing adapters were trimmed by using TrimGalore (length > 36bp and quality > 20) (Bolger et al. 2014). The clean data were mapped back to *Tetrahymena* MAC genome using HISAT2 with the parameters “--no-discordant” and “--no-spliced-alignment” (Anders et al. 2014). Reads with multiple mapping in the SAM file were deleted. Samtools (Li et al. 2009) was used to sort and remove potential PCR duplicates. Fragments in the range of 120bp-260bp were regarded as mononucleosomes and used for downstream analyses. The final results were calculated, normalized, and visualized by deepTools with the parameter “--MNase” in the bamCoverage sub-feature (Ramírez et al. 2014).

### Cross-linking ChIP-seq and data analysis

Approximately 5 × 10^7^ HA-tagged cells were fixed with 1% paraformaldehyde (Sigma, P6148) for 30 min. The fixation reaction was stopped by adding 25 mM Glycine. The fixed cells were resuspended in SDS lysis buffer (50 mM Tris-HCl pH 7.5, 5 mM EDTA, 1% SDS) and sonicated using a Bioruptor (Diagenode). After centrifugation, the supernatant was diluted 5 times with ChIP dilution buffer (167 mM NaCl, 16.7 mM Tris-HCl pH 8.1, 1.2 mM EDTA, 1.1% Triton X-100, 0.01% SDS) and immunoprecipitated with Pierce anti-HA magnetic beads (Thermo Scientific, 88836) at 4°C overnight.

Data trimming, mapping, PCR duplicates removal, and results display were processed using the same methods as in native ChIP-seq. Fragments in the range of 100bp-500bp were used for downstream analyses. Enriched genes were defined as ChIP/input ratio higher than 1 (normalized by total counts respectively).

### ATAC-seq and data analysis

Approximately 1 × 10^7^ purified MACs were treated with ATAC-resuspension buffer (10 mM Tris-HCl pH 7.5, 10 mM NaCl, 3 mM MgCl2, 0.1% NP40, 0.1% Tween20, 0.01% Digitonin) at 4°C for 3 min. MACs were resuspended in transposition buffer (10 mM Tris-HCl pH 7.5, 5 mM MgCl2, 10% dimethyl formamide, 0.1% Tween20, 0.01% Digitonin) and treated with Tn5 transposase (Active Motif, 81284) at 37°C for 30 min with 1000 rpm mixing.

The low quality paired-end reads of raw data were filtered and sequencing adapters were trimmed by using TrimGalore (length > 20 bp and quality > 20). Mapping, PCR duplicates removal, and results display were processed using the same methods as in native ChIP-seq. The ATAC-seq results were normalized by the background signal using customized Perl scripts, with analyses focused on the region from 200bp upstream to 100bp downstream of TSS.

## Acknowledgement

This work was supported by the National Natural Science Foundation of China (32125006) and Natural Science Foundation of Shandong Province of China (ZR2024ZD40). The authors would like to thank Dr. Jie Xiong (Institute of Hydrobiology, Chinese Academy of Sciences) for sharing the Con-A labeling protocol, Dr. Xiaoyuan Song for sharing the ATAC-seq protocol, and Dr. Chao Li, Mr. Liping Lv, Mr. Rui Wang, and Mr. Zhaorui Zhou (Ocean University of China, OUC) for providing their unpublished genomic data of multiple ciliates. High-performance computing resources for data processing were provided by the Institute of Evolution and Marine Biodiversity, High-Performance Biological Supercomputing Center, and Marine Big Data Center of Institute for Advanced Ocean Study at OUC. Our special thanks are given to Prof. Weibo Song (OUC) for his helpful suggestions during drafting the manuscript.

## Author contribution

J.N. H.Y. and S.G. conceived the study and designed the experiments; J.N. and H.Y. performed most of the experiments; L.N. constructed all mutant rescue strains. Y.L. constructed some knockout strains, B.N. constructed *DMT3*-CHA tagging strain, J.N. performed the bioinformatic analysis with H.Y., W.Z. and N.S. performed the mass spectrometry analysis; J.N. and S.G. wrote the paper, with inputs from all authors. S.G. conceived and supervised the project, wrote and revised the manuscript, and provided funding resources. All authors read and approved the final manuscript.

The authors declare no conflict of interest.

## Notes

### Competing Interest Statement

The authors have declared no competing interest.

